# Histone H3 availability is more important for development than H3.2 versus H3.3 subtype identity

**DOI:** 10.64898/2026.02.09.704946

**Authors:** Jeanne-Marie E. McPherson, Claire Sykes, Lucy C. Grossmann, Christina H. Hill, Mary P Leatham-Jensen, Robert J. Duronio, Daniel J. McKay

## Abstract

The distinct contributions of replication-dependent and replication-independent histones to development and genome function remain unclear. In this study, we investigate how the distinct protein identities of the histone H3.2 and H3.3 subtypes contribute to development and gene regulation in *Drosophila*. Comparing animals in which the replication-independent *H3.3* genes were mutated to produce the replication-dependent H3.2 protein with those carrying deletions of the replication-independent *H3.3* genes revealed that replication-independent H3.3 is essential for fertility, adult locomotor behavior, and normal longevity. However, development to adulthood does not depend on which replication-independent H3 subtype is expressed from the *H3.3* loci. Moreover, replication-independent H3.3 is not required to establish or maintain global patterns of chromatin accessibility or gene expression in the adult brain. Surprisingly, we find that expression of H3.2 from the replication-dependent *HisC* locus is essential in post-replicative cells in the absence of replication-independent H3.3, and we uncover a critical role for the HIRA histone chaperone complex in preserving genome function when replication-independent *H3.3* is deleted. We conclude that an available pool of H3 is more critical than the specific identity of H3 in the pool.

## INTRODUCTION

Histones are among the most abundant proteins in eukaryotic cells and play a fundamental role in organizing the genome. By packaging DNA into chromatin, histones control access to the information encoded in the genome, thereby regulating all DNA-templated processes, including transcription, replication, and repair. The basic structural unit of chromatin is the nucleosome, consisting of 147 base pairs of DNA wrapped around an octamer of histone proteins (Luger et al. 1997). This octamer is composed of four core histones – H2A, H2B, H3, and H4 – whose sequences are deeply conserved during evolution (Hocher and Warnecke 2024; Henikoff and Smith 2015). Because histone abundance directly influences chromatin structure, histone levels must be tightly controlled. Too few histones causes cell cycle arrest due to insufficient chromatin assembly (Günesdogan et al. 2014; Ye et al. 2003; Han et al. 1987), while excess histones are cytotoxic and promote genome instability (Singh et al. 2010; Maya Miles et al. 2018). Between these extremes, histone levels modulate nucleosome occupancy and positioning (Gossett and Lieb 2012; Mühlen et al.), thus shaping access to DNA regulatory elements like enhancers and promoters. Due to the profound impact of histone levels on genome function, it is essential to understand how cells maintain a steady and physiologically balanced histone supply.

Tight control of histone gene expression is achieved through both transcriptional and post-transcriptional mechanisms (Marzluff et al. 2008). At the transcriptional level, regulation differs between the two broad categories of histone genes: replication-dependent and replication-independent. Replication-dependent histone genes are expressed at high levels during S phase to meet the massive demand for new histones during DNA replication. By contrast, replication-independent histone genes are expressed at all cell cycle stages and in post-replicative cells to meet the demand for new histones that arises through nucleosome turnover. At the post-transcriptional level, excess histone proteins are targeted for proteosome-mediated degradation (Singh et al. 2009). Histone mRNA stability is likewise tightly regulated (Marzluff and Duronio 2002). For instance, replication-dependent histone mRNAs are degraded at the end of each S phase when the demand for histone proteins diminishes (Harris et al. 1991; Mullen and Marzluff 2008).

In addition to differing in the timing of their expression, many replication-independent histone genes encode “variant” proteins that differ in amino acid sequence from their replication-dependent counterparts (Martire and Banaszynski 2020; Henikoff and Smith 2015). These sequence differences mediate differential interactions with histone chaperone complexes and are thought to underlie specialized functions of histones in chromatin. In *Drosophila melanogaster*, histone H3.3 is the predominant replication-independent histone H3, whereas H3.2 is the only replication-dependent H3 subtype. The centromere-specific H3 variant *Cid* constitutes a small fraction of total H3 (Ahmad and Henikoff 2002a). H3.2 and H3.3 differ in their genomic distributions, with H3.2 enriched at many heterochromatic regions and H3.3 enriched at active genes and other sites of nucleosome remodeling (Mito et al. 2007). Nucleosomes that contain H3.3 and the H2A.Z histone variant have been proposed to be less stable than those that contain H3.2 (Jin et al. 2009; Jin and Felsenfeld 2007). H3.2 and H3.3 also differ in their post-translational modification (PTM) profiles, with H3.2 enriched in PTMs associated with gene repression like H3K27me3 and H3.3 enriched in acetylation and other PTMs associated with gene activation (McKittrick et al. 2004). H3.2 differs from H3.3 by just four amino acids. One of the four amino acid differences occurs at residue 31 in the N-terminal tail. Alanine 31 of H3.2 (H3.2A31) is non-modifiable, whereas serine 31 of H3.3 (H3.3S31) can be phosphorylated (H3.3S31ph) (Hake et al. 2005). H3.3S31ph has been proposed to stimulate activity of the p300 acetyltransferase and to participate in transcriptional induction in response to extracellular signals (Martire et al. 2019; Armache et al. 2020; Ma et al. 2025).

The remaining three amino acid differences between H3.2 and H3.3 are in the histone fold domain and modulate the interaction of each H3 subtype with chaperone complexes. Histone chaperones are a diverse class of proteins that work together to form an efficient histone supply chain, ensuring that nucleosomes are assembled when and where necessary (Hammond et al. 2017). During S phase, the massive influx of histones needed to package newly synthesized DNA into chromatin is managed by the H3.2-specific histone chaperone complex, CAF-1 (Rouillon et al. 2023; Verreault et al. 1996). Outside of S phase, the HIRA complex is responsible for depositing H3.3 at actively transcribed genes and regulatory regions independently of DNA replication (Banaszynski et al. 2013; Tang et al. 2012; Xiong et al. 2018). A different complex, DAXX/ATRX, mediates H3.3 loading at telomeres and pericentric heterochromatin (Ali Fromental-Ramain, 2017; Drané et al., 2010; Hoelper et al., 2017).

Despite the distinguishing features of H3.2 and H3.3, it has been difficult to determine whether proposed functional differences are due to differences in their amino acid sequence (“identity”) or differences in when they are synthesized and deposited into chromatin (“timing”). Most genetic tests of H3.3 function rely on deletion of *H3.3* genes. However, this approach changes both identity and timing by removing the H3.3 protein and its replication-independent expression, thus confounding interpretation. To address this problem, we used CRISPR to convert the two *H3.3* genes of *Drosophila* into versions that encode the H3.2 protein. In this way, we altered H3 identity while maintaining the replication-independent timing of H3 expression. We find that although both H3 identity and timing matter, an available pool of H3 is of paramount importance, regardless of the H3 identity within the pool. We further find that the consequences of *H3.3* deletion are made less severe due to *H3.2* gene expression. Moreover, we determine that cells are robust to changes in H3 identity in part through utilization of the replication-independent histone chaperone HIRA.

## RESULTS

### Generating replication-independent H3 mutants using CRISPR

To identify developmental and genomic contexts that specifically depend on H3.3 protein identity rather than replication-independent timing of H3 expression, we used CRISPR to create mutants that express the H3.2 protein from the endogenous *H3.3* genes. Like humans, *Drosophila* H3.3 is encoded by two genes, *H3.3A* and *H3.3B*, that produce identical proteins. We introduced nucleotide changes to convert each H3.3-specific amino acid to the H3.2 version (S31A, A87S, I89V, G90M) at both *H3.3* loci (**Fig. 1A**). Using long, single stranded DNA repair templates, we recovered 2 distinct protein-coding mutations at each locus: (1) conversion of all 4 amino acids, termed *H3.3^H3.2^*, and (2) conversion of only S31 to A31, termed *H3.3^S31A^*. These *H3.3A* and *H3.3B* alleles were then combined to create genotypes expressing a single type of replication-independent H3 protein. We also obtained new deletion alleles of each gene, termed *H3.3^Δ^* (**Fig. 1A,B**). We confirmed that the *H3.3^H3.2^*and *H3.3^S31A^* mutations do not impact steady-state *His3.3A* and *His3.3B* RNA levels, and that the *H3.3^Δ^* mutations result in the loss of full-length transcripts (**Fig. 1C-E**). Although the *H3.3A* deletion produces small amounts of truncated transcript, sanger sequencing revealed the presence of a frameshift, indicating it does not have H3 coding potential. Thus, from the onset of zygotic transcription, throughout development and into adulthood, *H3.3^H3.2^*and *H3.3^S31A^* mutants express replication-independent H3 genes with an altered protein identity, whereas *H3.3^Δ^* mutants lack any replication-independent H3 gene expression (**Fig. 1B**).

**Figure 1.**
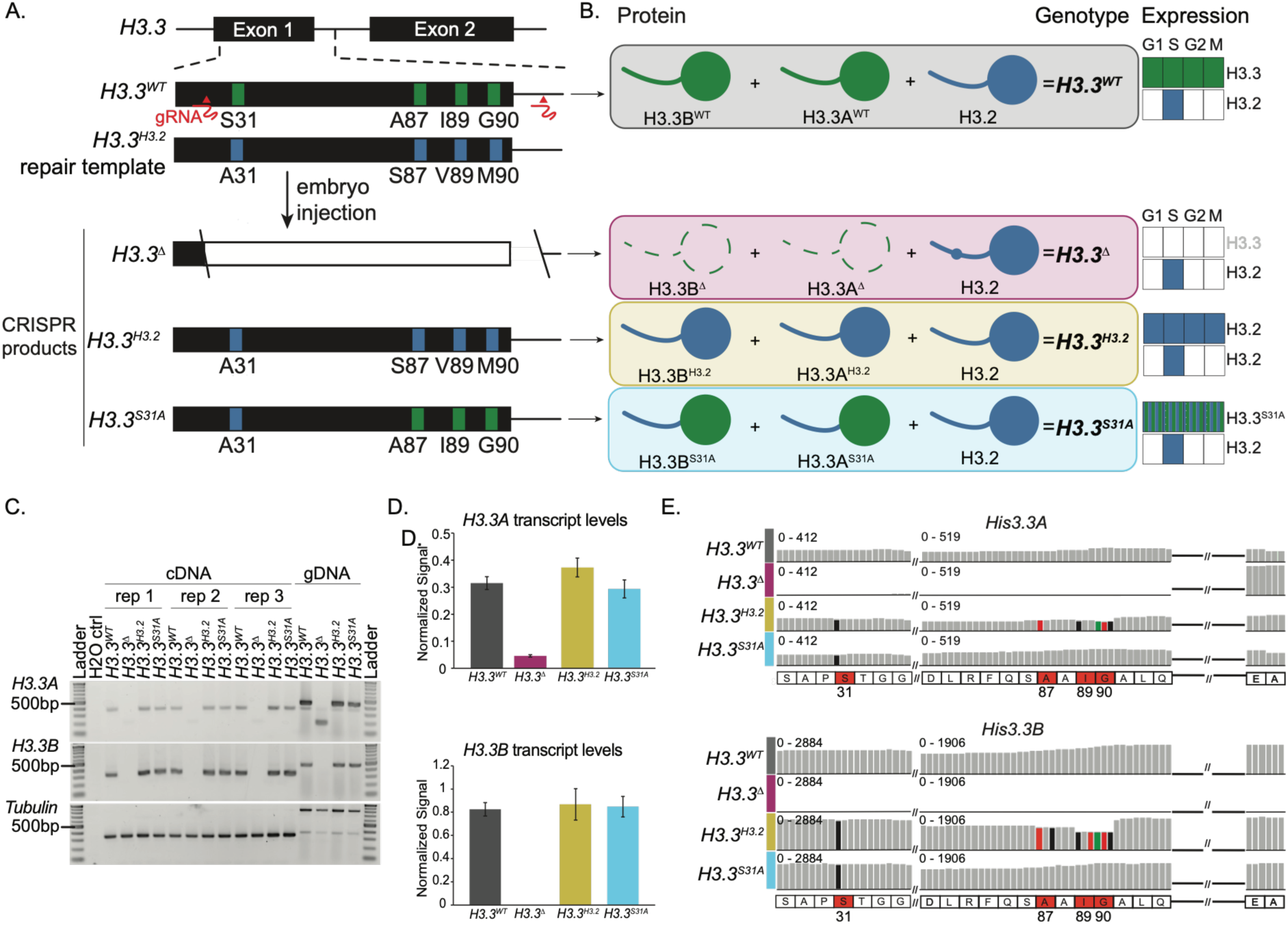
Generation of *Drosophila* genotypes expressing H3.2 from *H3.3* loci. (A) Schematic of the *H3.3A and H3.3B* gene structure and the long single-stranded DNA (lssDNA) repair template used for CRISPR/Cas9-mediated generation of mutant alleles (*H3.3^Δ^*, *H3.3^H3.2^*, and *H3.3^S31A^*). (B) Diagram of H3.2 and H3.3 protein present in each genotype and cell cycle timing of *H3.2* and *H3.3* gene expression. (C) Ethidium bromide-stained gel of RT-PCR products amplified from cDNA and gDNA templates for *H3.3A, H3.3B,* and *tubulin* from adults of the indicated genotypes. (D) Bar plot of tubulin normalized RT-PCR results for *H3.3A* (top) and *H3.3B* (bottom) in adults. Error bars: SEM for three biological replicates. (E) Browser shot of the *H3.3B* and *H3.3A* loci in 1d brain RNA-seq bam files for the indicated genotypes. Colored bars represent nucleotide changes (black = G, red = T, green = A) relative to WT (gray). Thick black line represents intron. Double slash represents an abbreviated sequence for display purposes.

### RI H3.3 is required for fertility but not for development to adulthood

We first sought to examine how eliminating replication-independent H3 gene expression, as compared to altering H3 subtype identity, affects viability and fertility. Deletion of both *H3.3A* and *H3.3B* (*H3.3^Δ^)* (**Fig. 2A**) resulted in a significant reduction in viability, with only 49.8% of expected females and 46.7% of expected males reaching adulthood (**Fig. 2A**). By contrast, *H3.3^H3.2^* and *H3.3^S31A^* mutants reached adulthood at expected frequencies **(Fig. 2A**). Full viability of *H3.3^H3.2^*and *H3.3^S31A^* indicates that a replication-independent supply of H3 contributes to normal development rather than the H3.3 protein itself. Consistent with prior findings (Sakai et al. 2009; Zhang et al. 2024), all *H3.3^Δ^* males were sterile (**Fig. 2B**). We observed that only 10% of *H3.3^Δ^* females produced eggs that hatched (**Fig. 2B**). The low fertility of *H3.3^Δ^* females is consistent with a prior report that *H3.3^Δ^* females have reduced fertility (Zhang et al. 2024) and with prior findings that H3.3 is required for unpacking sperm chromatin following fertilization (Loppin et al. 2005). Similar to *H3.3^Δ^*, we found that *H3.3^H3.2^* males are sterile and only 12.5% of *H3.3^H3.2^* females produce eggs that hatch (**Fig. 2B**). Unlike *H3.3^Δ^*, 15% of *H3.3^S31A^* males are fertile and 95% of *H3.3^S31A^* females produce eggs that hatch (**Fig. 2B**). These findings demonstrate that H3.3 is essential for male fertility and that H3.3S31 makes a critical contribution to this function. By contrast, H3.3 is required for female fertility independently of S31. Overall, we conclude that fly development to adulthood is insensitive to the identity of replication-independent H3 subtype whereas replication-independent H3.3 is critical for reproductive function.

**Figure 2.**
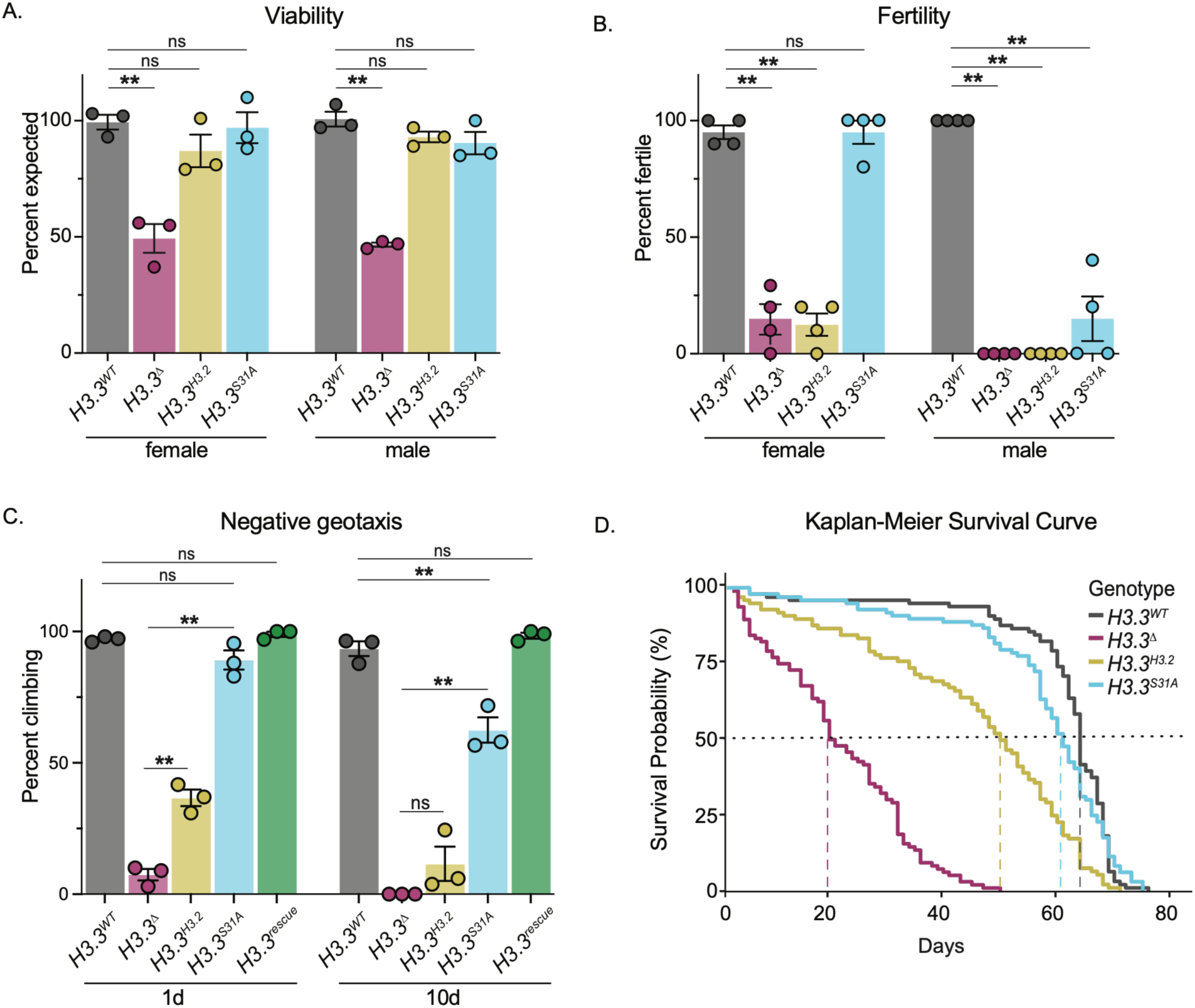
Replication-independent H3.3 is not required for development to adulthood, but is required for adult fertility, locomotor behavior, and longevity. (A) Bar plot of female and male viability for the indicated genotypes. Percent of expected genotypic frequencies based on Mendelian ratios. Dots represent individual biological replicates. B) Bar plot of fertility assay quantifying the percentage of females and males that produce eggs that hatch for the indicated genotypes. Each dot represents the mean of 10 animals tested per biological replicate. (C) Bar plot of negative geotaxis assay assessing climbing ability at 1 day (1d) and 10 days (10d) post-eclosion for the indicated genotypes. Each dot represents the mean of 10 animals tested per biological replicate. (D) Kaplan-Meier survival curves comparing longevity among the indicated genotypes. Dashed vertical lines indicate median lifespan for each genotype. Dotted horizontal line indicates 50% survival. H3.3^Δ^ and *H3.3^H3.2^* have a statistically significant reduction in longevity compared to *H3.3^WT^* by log-rank test. C, chi-square test (** p ≤ 0.01); D-E, One-way ANOVA with post-hoc Tukey HSD (** p ≤ 0.01). A-C, error bars: SEM for three biological replicates.

### Replication-independent H3.3 is essential for adult locomotion and longevity

Replication-independent histones are the primary source of new histones in non-replicating cells. As a result, H3.3 increases in abundance in post-mitotic cells and becomes the predominant H3 subtype in chromatin during aging (Funk et al. 2022; Pantazis and Bonner 1984; Urban and Zweidler 1983; Maze et al. 2015). Due to this accumulation, we reasoned that H3 identity might be particularly important in adulthood. We therefore examined whether *H3.3^Δ^*, *H3.3^H3.2^,* and *H3.3^S31A^* mutants exhibit normal behavior after reaching adulthood. We assessed adult locomotor function using negative geotaxis, a natural climbing response in *Drosophila* that declines with age (Rhodenizer et al. 2008a). At 1-day post-eclosion (1d), only 7.3% of *H3.3^Δ^*mutants successfully completed a climbing test compared to 97.1% of *H3.3^WT^*controls (**Fig. 2C**). By 10 days (10d), 0% of *H3.3^Δ^* mutants completed the climbing test compared to 93.8% of *H3.3^WT^* controls (**Fig. 2C**). To distinguish between the effects of loss of H3.3 protein identity from loss of replication-independent H3 expression, we evaluated climbing in *H3.3^H3.2^*and *H3.3^S31A^* mutants. *H3.3^H3.2^* mutants partially rescued the severe climbing defect observed in *H3.3^Δ^* mutants at 1d, with 36.5% climbing, but they were similarly defective to *H3.3^Δ^* mutants by 10d, with only 11.4% climbing (**Fig. 2C**). *H3.3^S31A^* mutants fully rescued the *H3.3^Δ^*climbing defect at 1d (88.8% climbing) but exhibited age-related decline by 10d (62.2% climbing) (**Fig. 2C**). Introducing a *H3.3B^WT^* transgene into *H3.3^H3.2^*mutants (*H3.3^rescue^*) fully restored climbing ability, indicating that negative geotaxis requires H3.3 protein (**Fig. 2C**). Collectively, these data indicate that H3.3 protein identity is necessary for normal locomotor behavior, and that replication-independent expression of H3.2 from the *H3.3* loci cannot fully substitute for loss of H3.3. Furthermore, the milder phenotype observed in *H3.3^S31A^*mutants compared to *H3.3^H3.2^* suggests that the three amino acid differences in the H3.3 globular domain contribute significantly to locomotor function.

Because negative geotaxis declines with aging (Rhodenizer et al. 2008b), we next examined longevity. We observed that *H3.3^Δ^* flies have a median longevity of 20 days, which is significantly shorter than the 64 days in *H3.3^WT^* controls (**Fig. 2D**). *H3.3^H3.2^* mutants also exhibited a significantly reduced longevity (median 50 days), but this reduction in longevity is not as significant as that of *H3.3^Δ^*, indicating that replication-independent timing of H3 expression contributes more to longevity than H3 subtype identity (**Fig. 2D)**. By contrast, *H3.3^S31A^* mutants had no significant difference in longevity relative to *H3.3^WT^*controls (median 62 days) (**Fig. 2D**). These findings parallel the locomotor data and demonstrate that although replication-independent expression of H3.2 can partially rescue the absence of H3.3, the globular domain of H3.3 is necessary for normal adult longevity in *Drosophila*.

### Adult brain chromatin accessibility is robust to loss of H3.3

Given the strong enrichment of H3.3 at *cis*-regulatory elements and nuclease-hypersensitive sites (Mito et al. 2007) and that both *H3.3^Δ^* and *H3.3^H3.2^* mutants have locomotor and longevity defects, we hypothesized that H3.3 may be critical for shaping the gene regulatory landscape of post-replicative cells. To test this hypothesis, we examined chromatin accessibility by performing ATAC-seq in *H3.3^WT^, H3.3^Δ^*, *H3.3^H3.2^*, and *H3.3^S31A^* adult brain, which is composed nearly entirely of post-replicative cells (Fogarty et al. 2024; Siegrist et al. 2010), at 1d and 10d after eclosion. Across all genotypes and the two time points, we identified a union set of 11,145 ATAC-seq peaks. First, we assessed whether temporal changes in chromatin accessibility could be identified in *H3.3^WT^* brains. Of the 11,145 peaks, only 19 (0.17%) exhibited significant differential accessibility between 1d and 10 days after eclosion (2 gained and 17 lost), indicating there is very little change in chromatin accessibility in wild-type aging brains during the first 10 days of adult life (**Fig. 3A**). In addition, these changes required neither H3.3 protein identity nor replication-independent timing of H3 gene expression, as we found that all 19 showed comparable log_2_ fold changes from 1d to 10d after eclosion in *H3.3^Δ^*, *H3.3^H3.2^*, and *H3.3^S31A^* mutant brains (**Fig. 3B,C**). Likewise, pairwise-comparisons of the entire ATAC-seq dataset between 1d and 10d for each *H3.3* mutant genotype revealed similarly subtle changes in chromatin accessibility (**Supplemental Fig. S1A-C).**

**Figure 3.**
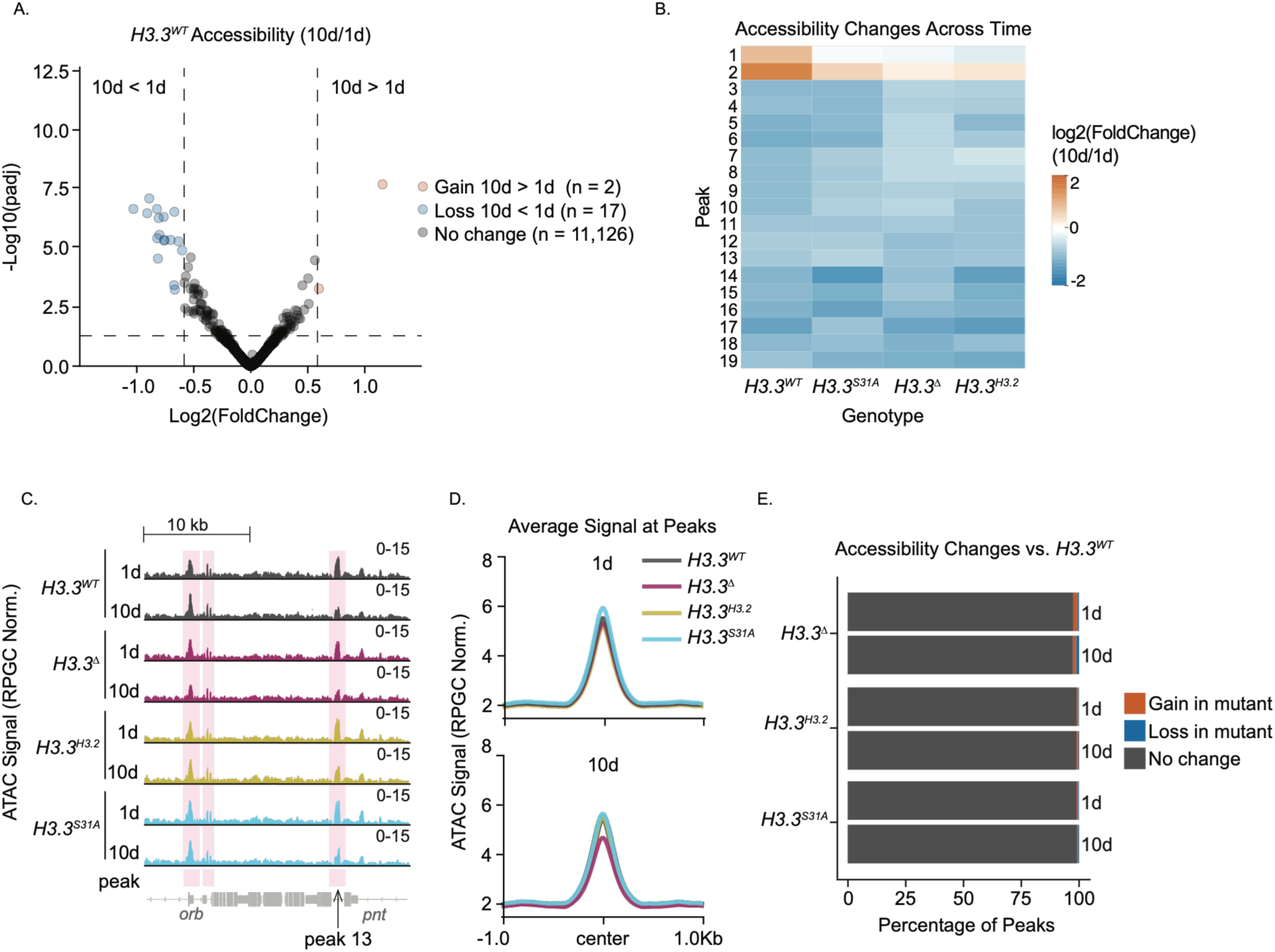
Adult brain chromatin accessibility is robust to loss of H3.3. (A) Volcano plot of ATAC-seq data comparing 1-day (1d) and 10-day (10d) old brain tissue in *H3.3^WT^*. Differentially accessible regions (P_adj_ < 0.05, absolute log_2_ fold change > 0.585) are colored orange and blue. Peaks are categorized as gained (n=2), lost (n = 17), or not changed (n= 11,126). (B) Heatmap of log_2_ fold changes in peak accessibility across the indicated genotypes from 1d to 10d. (C) Browser shot of RPGC-normalized ATAC signal from *H3.3^WT^* (gray), *H3.3^Δ^*(magenta), *H3.3^H3.2^* (yellow), and *H3.3^S31A^* (blue) brains at 1d and 10d. Pink highlights correspond to temporally dynamic (peak 13) and static peaks. (D) ATAC average signal plots centered on peak summit (±1 kb) for each genotype at 1d and 10d. (E) Bar plot showing the percentage of peaks with accessibility changes relative to *H3.3^WT^* at 1d and 10d. Peaks are categorized as gained (orange), lost (blue), or not changed (gray).

Second, we performed pair-wise comparisons of each mutant to *H3.3^WT^*controls at both 1d and 10d. In all instances, chromatin accessibility profiles remain largely unchanged (**Fig. 3D,E**). For example, only 3% of ATAC peaks were differentially accessible in *H3.3^Δ^* mutants compared to *H3.3^WT^* at 1d (**Fig. 3E**), consistent with prior reports of subtle chromatin accessibility defects in *H3.3* knockout mouse embryonic stem cells (Martire et al. 2019; Tafessu et al. 2023). Even fewer ATAC peaks were identified as differentially accessible in *H3.3^H3.2^* and *H3.3^S31A^* 1d brains (**Fig. 3E**). Moreover, chromatin accessibility profiles in each of the *H3.3* mutants were unaffected by aging, with only 2.5%, 1.0%, and 1.1% of ATAC peaks identified as differentially accessible in *H3.3^Δ^*, *H3.3^H3.2^*, and *H3.3^S31A^* 10d old brains, respectively (**Fig. 3E**). These data reveal that neither replication-independent timing of H3 gene expression nor H3.3 protein identity is required to establish or maintain chromatin accessibility patterns in the brains of adult flies through 10d after eclosion. The lack of chromatin accessibility changes in *H3.3^H3.2^*mutants suggest that, in the absence of H3.3, H3.2 may be deposited into chromatin similarly to H3.3 in brains.

### The adult brain transcriptome is robust to loss of H3.3

H3.3 incorporation into chromatin has been proposed to impact transcription factor binding (Tafessu et al. 2023). Therefore, although we observed no large-scale defects in chromatin accessibility profiles in H3.3 mutants, we next used total RNA sequencing to examine steady-state mRNA levels in the brains of *H3.3^WT^*, *H3.3^Δ^*, *H3.3^H3.2^*, and *H3.3^S31A^*animals at 1d and 10d after eclosion. Examination of RNA-seq data confirmed the intended mutations were present at both *H3.3A* and *H3.3B* loci in all genotypes (**Fig. 1E**). We asked how the wild-type adult brain transcriptome changes from 1d to 10d. Among the 12,557 expressed genes in *H3.3^WT^* animals, 447 (3.56%) showed significant changes in expression from 1d to 10d (**Fig. 4A**). GSEA revealed that these genes are associated with biological pathways known to be affected by aging in the brain, such as increased ciliary function and decreased cellular respiration and synaptic signaling (**Supplemental Fig. S2A**) (Ferguson et al. 2005; Petralia et al. 2014; Youn and Han 2018). Remarkably, 406 of these 447 genes (90.8%) exhibit similar directional changes in expression in *H3.3^Δ^*, *H3.3^H3.2^*, and *H3.3^S31A^* mutants (**Fig. 4B**), and GSEA identified many of the same biological pathways in these mutants (**Supplemental Fig. S2B-D**).

**Figure 4.**
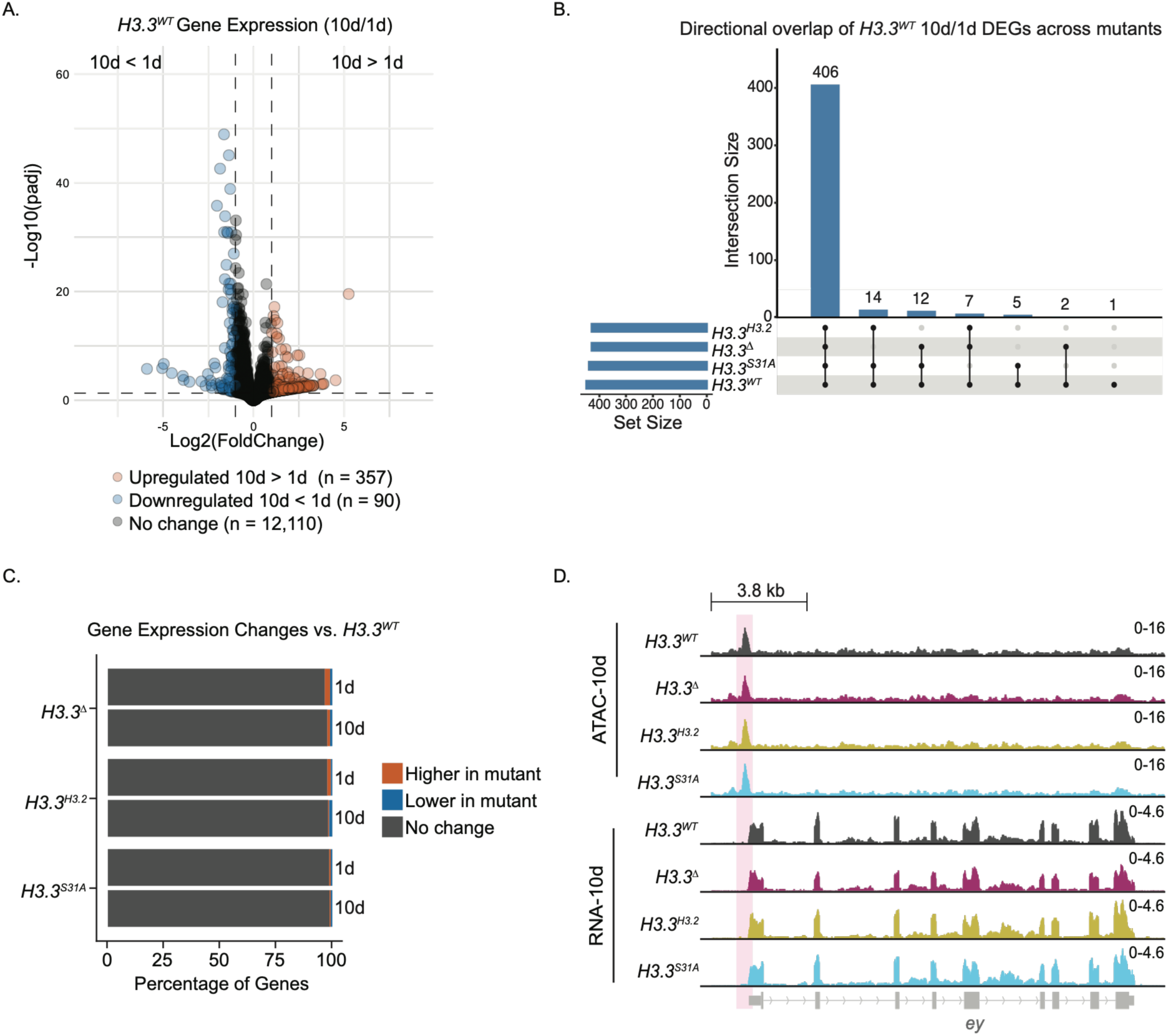
The adult brain transcriptome is robust to loss of H3.3. (D) Volcano plot of RNA-seq data comparing 1-day (1d) and 10-day (10d) old brain tissue in *H3.3^WT^*. Differentially expressed genes (P_adj_ < 0.05, absolute log_2_ fold change > 1.0) are colored orange and blue. Genes are categorized as upregulated (orange), downregulated (blue), or not changed (gray). (E) UpSet plot showing overlap among genes that are differentially expressed in 1d vs. 10d old *H3.3^WT^*brains (n = 447) and exhibit concordant directionality of expression change in each H3.3 mutant (*H3.3^S31A^, H3.3^Δ^,* and *H3.3^H3.2^*) across time. Bars indicate the number of genes shared among the indicated sets, and intersections represent genes with consistent directional regulation across *H3.3^WT^* and one or more mutant genotypes.

Next, we examined how gene expression at 1d and 10d is impacted in *H3.3^Δ^*, *H3.3^H3.2^*, and *H3.3^S31A^*mutants through direct pair-wise comparisons with *H3.3^WT^* controls. At 1d, we identified 376 differentially expressed genes (DEGs) in *H3.3^Δ^*brains (2.83% of expressed genes) (**Fig. 4C**). Similar numbers of DEGs were identified in *H3.3^H3.2^* (249 DEGs, 1.88% of expressed genes) and *H3.3^S31A^* (148 DEGs, 1.12% of expressed genes) mutants (**Fig. 4C**), indicating modest transcriptional changes across all genotypes. We observed similar small differences in gene expression at 10d in *H3.3^Δ^* (242 DEGs, 1.82% of expressed genes), *H3.3^H3.2^* (224 DEGs, 1.69% of expressed genes), and *H3.3^S31A^* (120 DEGs, 0.9% of expressed genes) mutants (**Fig. 4C**). Supporting these observations, chromatin accessibility and transcript abundance at the *eyeless* (*ey*) locus, a gene expressed in neurons, were similar across all genotypes at 10d (**Fig. 4D)**. These data indicate that steady-state gene expression in aging brains does not require H3.3 protein identity and is largely independent of replication-independent H3 expression. Together with our observations from chromatin accessibility profiling, these findings indicate that cells have robust compensatory mechanisms that buffer against changes in histone H3 supply which act to preserve the gene regulatory landscape.

### A homeostatic mechanism maintains total H3 levels when replication-independent H3 identity is altered

How might cells buffer against changes in histone H3 supply? Mechanisms that sense and respond to changes in replication-independent timing and H3 subtype identity could act at multiple levels including transcription, translation, histone protein turnover, or histone protein recycling. We considered the possibility that in the absence of H3.3, expression of *H3.2* from the *HisC* locus is increased to provide a pool of H3 in post-replicative cells. To examine this possibility, we assessed histone gene expression in our RNA-seq datasets. We first compared normalized steady-state levels of wild-type replication-dependent histone mRNAs in post-replicative 1d and 10d old brains to proliferating cells from third larval instar wing imaginal discs (Stutzman et al. 2024). We found that the replication-dependent *H3.2* genes were expressed in *H3.3^WT^* brains, but that their transcript levels were low compared to wing imaginal discs (93% and 73% decrease in 1d and 10d, respectively) (**Fig. 5A,B**). The other replication-dependent histone mRNAs behaved similarly to H3.2, although the magnitude of the reduction relative to wing imaginal discs was not as great as H3.2 (**Fig. 5A,B**). Surprisingly, in 10d brains we found evidence for transcripts extending beyond the normal replication-dependent histone mRNA 3’ ends (**Fig. 5A**), which in proliferating cells are not polyadenylated (Marzluff and Koreski 2017). This observation is consistent with the use of cryptic downstream polyadenylation signals as occurs when components of the replication-dependent histone mRNA 3’ end processing machinery are mutated (Lanzotti et al. 2002). Polyadenylation of replication-dependent histone mRNA has also previously been observed in wild-type post-replicative mammalian cells (Lyons et al. 2016). This alternative mechanism for generating mature replication-dependent histone mRNA could explain the relative increase in *H3.2* transcripts in 10d versus 1d brains (**Fig. 5A,B**). These data demonstrate that replication-dependent histone genes are normally expressed in post-mitotic brain cells, albeit at much lower levels than in proliferating cells, potentially contributing to the H3 protein supply, and that their transcripts are processed differently in aged brains.

**Figure 5.**
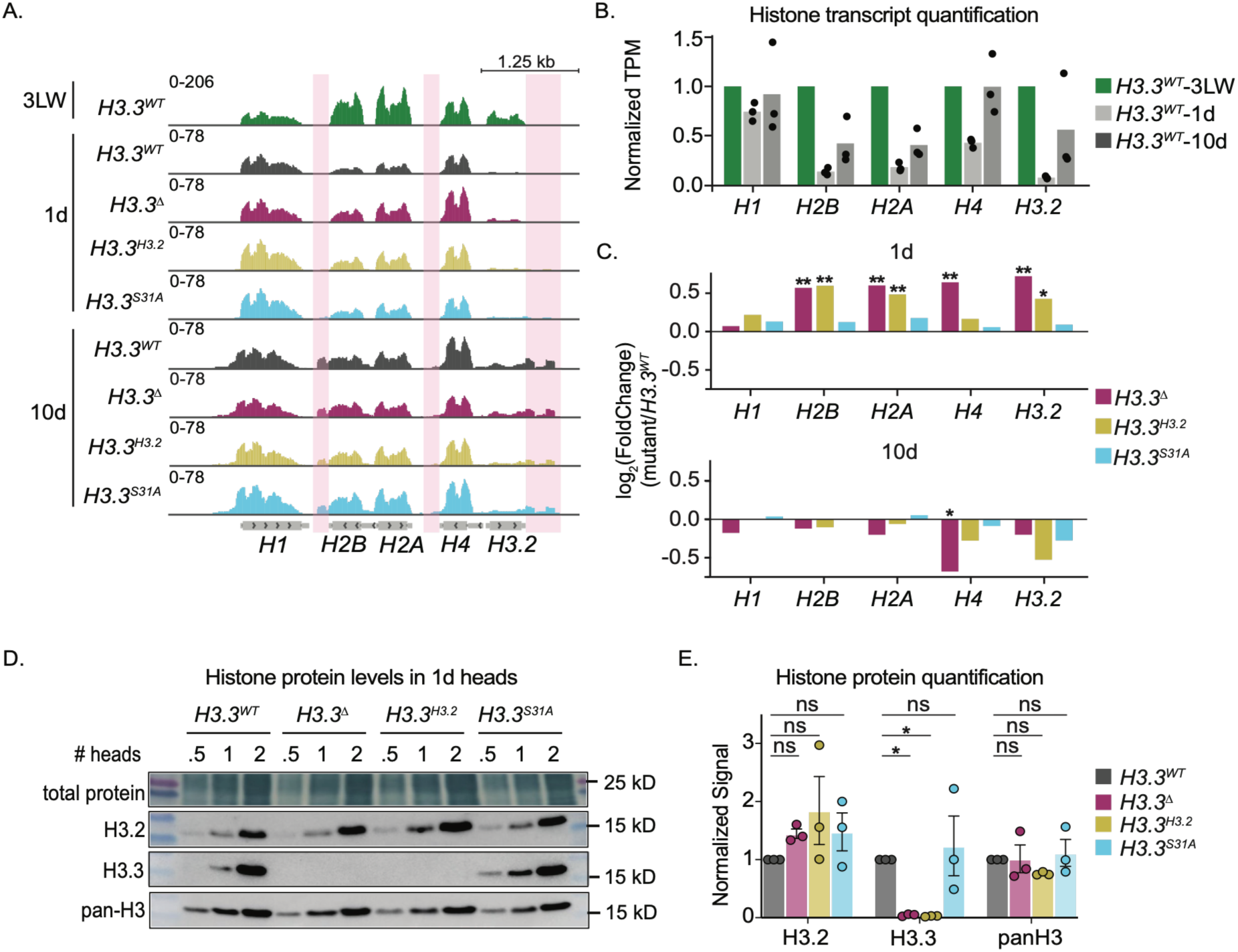
A homeostatic mechanism maintains total H3 levels when replication-independent H3 identity is altered. (A) Browser shot of RPGC normalized RNA signal at the replication-dependent histone locus in third larval instar wing imaginal discs (3LW), 1-day (1d) old brains, and 10-day old (10d) brains for the indicated genotypes. The locations of reads associated with polyadenylated replication-dependent histone transcripts are highlighted in pink. Signal indicates pooled replicates; we excluded *H3.3^WT^*-10d-rep2 in the browser for visualization purposes due to high variance; inclusion of this replicate did not affect differential expression calls. (B) Bar plot of fraction of transcripts per million (TPM) normalized reads for the replication-dependent histone genes in each indicated genotype relative to *H3.3^WT^*-3LW mean TPM. Dots represent biological replicates. (C) Bar plots representing the log_2_ fold change in replication-dependent histone transcript levels in the brains of the indicated genotypes relative to *H3.3^WT^* at 1d (top) and 10d (bottom). Bars represent the mean of biological replicates. Statistical significance by DESeq2 Wald test with Benjamini–Hochberg adjusted p-values (* p ≤ 0.05 and ** p ≤ 0.01). (D) Western blot for H3.2, H3.3, and pan-H3 from 1d heads of the indicated genotypes. Dilution series with indicated number of heads for each genotype. (E) Bar plot depicting the mean fold change in H3.3, H3.2, and pan-H3 signal in the indicated genotypes relative to *H3.3^WT^* signal normalized to total protein. Error bars: SEM for three biological replicates (dots). Statistical significance was assessed using a one-way ANOVA followed by Dunnett’s post-hoc test (* p ≤ 0.05).

To determine whether replication-dependent H3.2 mRNA levels increase in the absence of H3.3, we compared replication-dependent histone mRNA levels in *H3.3^Δ^*, *H3.3^H3.2^*, and *H3.3^S31A^* mutants to *H3.3^WT^* controls. We observed that *H3.2* mRNA levels were elevated in 1d *H3.3^Δ^*brains compared to *H3.3^WT^*, as were the *H2A*, *H2B*, and *H4* mRNAs (**Fig. 5C**). Similarly, *H3.2*, *H2A*, and *H2B* mRNA levels were elevated in *H3.3^H3.2^* mutants relative to *H3.3^WT^* (**Fig. 5C**). By contrast, no replication-dependent histone mRNAs were elevated in *H3.3^S31A^* brains (**Fig. 5C**). These data suggest that the replication-dependent *H3.2* genes may compensate for the absence of H3 expression from the replication-independent H3.3 loci in 1d old brains. By 10d, replication-dependent histone gene expression in all mutants largely normalized to *H3.3^WT^* levels (**Fig. 5C**). An exception was *H4* mRNA, which was significantly lower in *H3.3^Δ^* mutants at 10d (**Fig. 5C**), perhaps to match reduced H3 availability.

To assess whether H3 protein levels change in the absence of H3.3, we measured total H3 abundance in the heads of 1d old animals using a pan-H3 antibody that recognizes both H3.2 and H3.3. Notably, total H3 protein levels were comparable across all genotypes, including *H3.3^Δ^*(**Fig. 5D,E**). We also measured H3.3 and H3.2 protein levels in 1d old heads using antibodies specific to each H3 subtype. As expected, H3.3 protein is absent in *H3.3^Δ^*and *H3.3^H3.2^* mutants but present at levels comparable to *H3.3^WT^*control in the *H3.3^S31A^* mutant, consistent with the antibody’s recognition of the AIG motif in the H3.3 globular domain, which remains intact in *H3.3^S31A^* (**Fig. 5D,E**). We observed a 1.4-fold increase in H3.2 protein in the *H3.3^Δ^* mutant compared to *H3.3^WT^*. Although this increase did not reach statistical significance, it is consistent with elevated *H3.2* transcripts in 1d brains (**Fig. 5D,E**). Likewise, *H3.3^H3.2^* mutants exhibited a 1.8-fold increase in H3.2 protein levels compared to *H3.3^WT^*controls (**Fig. 5D,E**), consistent with production of H3.2 protein both from the replication-dependent *H3.2* genes and from the replication-independent *H3.3* loci. Although the epitope recognized by the H3.2 antibody is proprietary, the elevated H3.2 signal in *H3.3^S31A^* mutants suggests that it likely contains H3.2A31. Altogether, quantification of RNA and protein levels suggests that in the absence of replication-independent H3.3, a homeostatic mechanism engages to maintain a pool of H3 in post-replicative brain cells by increasing replication-dependent H3.2 expression, consistent with a previous observation that total H3 levels are unchanged in H3.3-knockout mouse neurons (Funk et al. 2022).

### Replication-dependent H3.2 is important in the absence of replication-independent H3

Expression of H3.2 from the *H3.3* loci rescues the viability defect observed in *H3.3^Δ^*genotypes, indicating that replication-independent expression of histone H3 *per se* is more important for completion of development than whether that replication-independent H3 protein is H3.3. A corollary to this conclusion is that cells can use H3.2 outside of S phase in the absence of replication-independent expression of H3.3. Interestingly, our RNA-seq analyses revealed that H3.2 mRNAs are normally present in wild-type adult brains, and *H3.*3*^Δ^* animals show signs of increased H3.2, raising the possibility that H3.2 expression reduces the severity of loss of H3.3 in post-mitotic cells. We therefore tested whether H3.2 expression compensates for loss of H3.3 through two lines of investigation. First, we genetically reduced the copy number of all replication-dependent histone genes from the wild-type 200 copies to 12 (*12x^HWT^*). We found that whereas *H3.3^WT^*; *12x^HWT^* animals are viable at expected frequency, *H3.3^H3.2^; 12x^HWT^* animals are completely inviable (**Fig. 6A**). These data indicate that H3.3 protein identity is required for viability when replication-dependent histone gene copy number is reduced, extending our prior findings with *H3.3^Δ^; 12xHWT* animals (McPherson et al. 2023). In an orthogonal approach, we knocked down H3.2 specifically in post-mitotic cells using shRNAs targeting the H3.2 3’UTR (Arroyo et al. 2025). shRNA expression in mature neurons using *nSyb-GAL4* did not affect the viability of wild-type or *H3.3^H3.^*^2^ animals (**Fig. 6B**), indicating that endogenous H3.2 is not required when H3 is expressed outside of S phase. By contrast, shRNA expression did result in a significant reduction in viability of H3.3*^Δ^* animals, with only 24.1% of expected eclosing (**Fig. 6B**, McPherson et al. 2023). Altogether, we conclude that replication-independent H3.2 expression can support viability of post-replicative cells. Moreover, endogenous H3.2 becomes essential in post-mitotic cells in the absence of replication-independent H3 expression.

**Figure 6.**
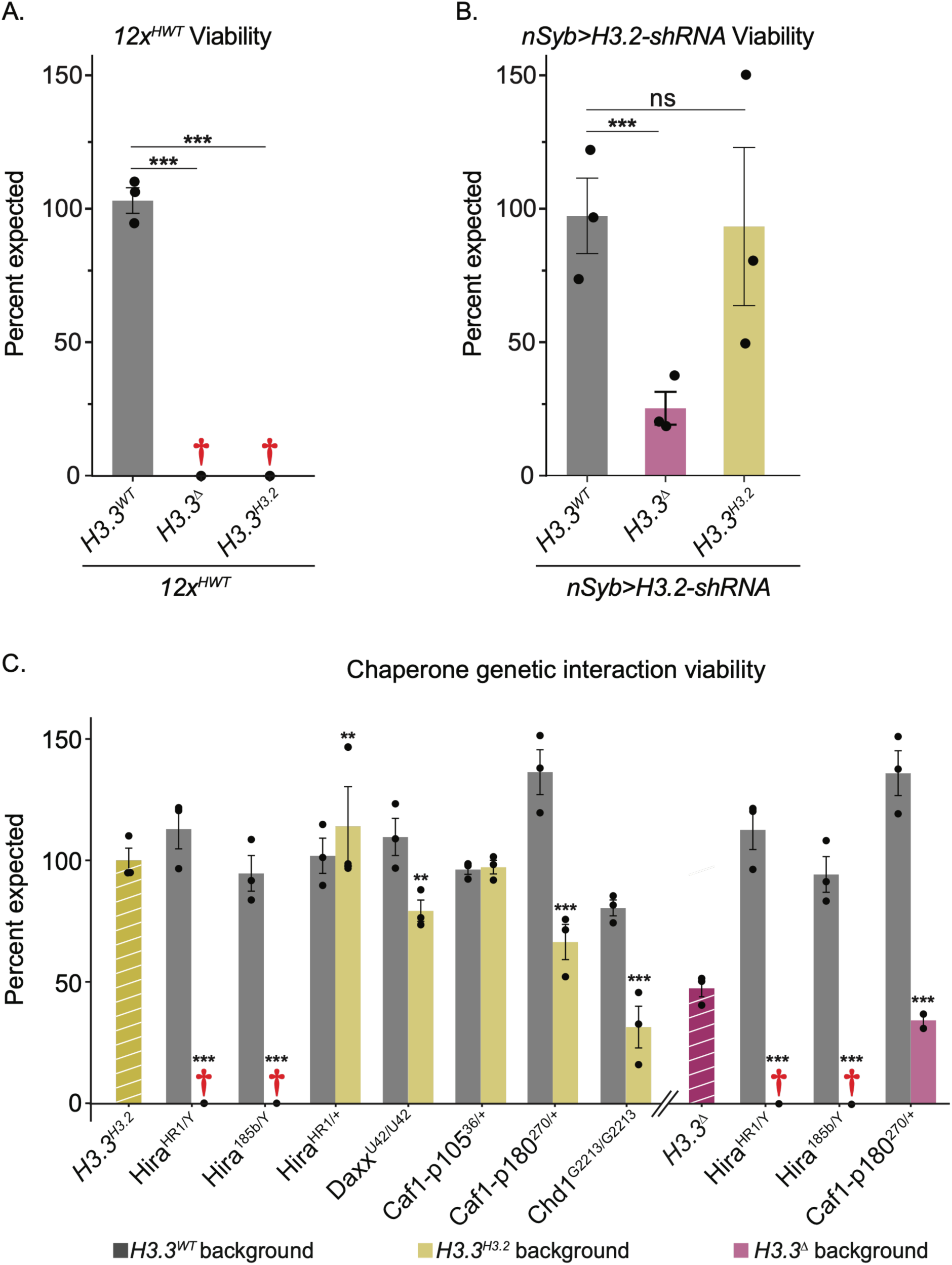
Hira is required for development in the absence of replication-independent H3.3. (A) Bar plot of viability of *H3.3^WT^*, *H3.3^Δ^*, and *H3.3^H3.2^* animals with reduced replication-dependent histone gene copy number (*12x^HWT^*). (B) Bar plot of viability of *H3.3^WT^*, *H3.3^Δ^*, and *H3.3^H3.2^* animals expressing *nSyb-Gal4>UAS-H3.2-shRNA*. (C) Bar plot of viability of *H3.3^WT^* (gray), *H3.3^H3.2^*(yellow) and *H3.3^Δ^* (magenta) animals with the indicated chaperone mutations. Hashed bars represent control genotypes. A-C, error bars: SEM for three biological replicates (dots). Asterisks indicate statistical significance by Chi-square test (* p <0.05, ** p < 0.01, *** p <0.0001, † dead).

### The histone chaperone Hira is required in the absence of replication-independent H3.3

Our finding that H3.2 compensates for the loss of replication-independent H3.3 suggests a model in which a minimum pool of H3 must be maintained to preserve genome function regardless of H3 subtype identity. Because histone chaperones are essential for regulating the flux of histones into chromatin, they may play a central role in this homeostatic mechanism by sensing and responding to changes in the available histone pool to maintain sufficient levels of H3 in chromatin. Consistent with this idea, depletion of CAF-1 in breast cancer cells leads to upregulation of HIRA (Gomes et al. 2019), and in certain ALT cancers, loss of ATRX causes redistribution of HIRA to genomic regions typically enriched for ATRX (Hoang et al. 2020). We therefore hypothesized that one or more histone chaperone complexes were necessary for genome function upon changing replication-independent H3 identity. We reasoned that reduction or loss of a histone chaperone needed to deposit replication-independent H3.2 protein into the genome may reduce the viability and/or exacerbate the poor locomotion of *H3.3^H3.2^* mutants. We tested both members of the H3.2-specific chaperone complex CAF-1 and members of the H3.3-specific chaperone complexes HIRA and DAXX/ATRX. We also tested Chd1, a chromatin remodeler involved in H3.3 deposition in the brain (Schoberleitner et al. 2021).

We found that removing one copy of the *Caf1-p105* gene in *H3.3^H3.2^* mutants (*Caf1-p105^36/+^*) did not significantly impact viability or adult locomotion (**Fig. 6C, Supplemental Fig. S3**). Heterozygosity of *Caf1-p180* in *H3.3^H3.2^* mutants *(Caf1-p180^270/+^)* caused a moderate, but significant, reduction in viability compared to control *H3.3^H3.2^* animals *(Caf1-p180^+/+^),* with 66.4% of expected surviving to adulthood (**Fig. 6C**). *Caf1-p180^270/+^*did not impact the performance of *H3.3^H3.2^* mutants in the climbing assay (**Supplemental Fig. S3**). Homozygous deletion of either *Caf1-p105* or *Caf1-p180* is lethal and thus these genotypes could not be examined. Homozygous deletion of *Daxx* in *H3.3^H3.2^* mutants (*Daxx^U42/U42^*) caused a modest reduction in viability, with 79.3% of expected surviving to adulthood, but *Daxx* removal did not impact the *H3.3^H3.2^* climbing phenotype (**Fig. 6C, Supplemental Fig. S3**). Similarly, homozygous deletion of *Chd1 (Chd1^G2213/G2213^)* caused a significant reduction in *H3.3^H3.2^* viability, with 31.7% of expected surviving to adulthood, but its deletion did not impact the *H3.2^H3.2^* climbing phenotype (**Fig. 5C, Supplemental Fig. S3).** Strikingly, two independent *Hira* loss-of-function alleles (*Hira^HR1/Y^*, *Hira^185b/Y^*) caused complete lethality of *H3.3^H3.2^* males, whereas *Hira* mutation had no impact on viability of *H3.3^WT^* controls (**Fig. 6C)**. These results indicate that HIRA function becomes essential when H3.2 is the source of replication-independent H3. We observed similar outcomes in *H3.3^Δ^* mutants, wherein *Hira* loss resulted in complete lethality (**Fig. 6C**). We conclude that the HIRA histone chaperone complex is required for development to adulthood in the absence of replication-independent H3.3. The lethality observed in both *H3.3^H3.2^* and *H3.3^Δ^*mutants suggests that, when H3.3 is absent, the HIRA complex engages H3.2 to deposit H3 into chromatin to maintain genome function during development.

## DISCUSSION

### Defining H3.3-specific functions in development and adult physiology

Regulation of histone expression and function establishes the chromatin landscapes necessary for maintaining cellular and organismal function and health. Dysregulation of these processes contributes to aging and a wide spectrum of pathologies, including cancer and neurodevelopmental disorders (Amodeo et al. 2025; Mossink et al. 2021; Wang et al. 2022). Despite its importance, the relative contributions of replication-dependent histones versus replication-independent histones to organism development and function remain incompletely understood. Previous studies in *Drosophila* demonstrated that mutating *H3.3* genes reduces viability and causes male sterility (Hödl and Basler 2009; McPherson et al. 2023; Sakai et al. 2009), but whether these phenotypes resulted from loss of H3.3 protein identity or the absence of replication-independent H3 expression was unclear. In this study, we engineered the *Drosophila* genome using CRISPR to encode H3.2 from both endogenous *H3.3* loci to alter H3 subtype identity while preserving the replication-independent timing of H3 expression. Our analysis revealed that *H3.3^H3.2^* mutants do not suffer from the reduced viability observed in *H3.3^Δ^* mutants, indicating that replication-independent timing of H3 expression is more important for the completion of development than any unique chromatin properties provided by H3.3 identity. Thus, a remarkable amount of cell-type specification and differentiation in *Drosophila* occurs in the absence of H3.3. Despite completing development normally, *H3.3^H3.2^* adult flies display several defects, including infertility, improper negative geotaxis, and reduced longevity. The negative geotaxis and longevity defects in *H3.3^H3.2^* mutants were less severe than in *H3.3^Δ^* mutants, indicating that H3.2 can partially compensate for loss of replication-independent H3, but it does not substitute for every H3.3 function.

### H3.3S31 is largely dispensable in Drosophila, except in the male germ line

Recent studies show that H3.3S31ph promotes signal-dependent transcription in mouse macrophages by reshaping the H3.3 PTM landscape. Specifically, H3.3S31ph blocks ZMYND11 binding to H3.3K36me3 while enhancing SETD2-mediated H3K36 methylation (Armache et al. 2020). H3.3S31ph has also been shown to facilitate DNA accessibility both by stimulating p300 acetyltransferase activity, which promotes acetylation at enhancers, and by enhancing FACT complex recruitment. (Martire et al. 2019; Ma et al. 2025; Sitbon et al. 2020). We found that *H3.3^S31A^* animals develop normally to adulthood and lack the locomotion and longevity defects of *H3.3^H3.2^* animals, suggesting these phenotypes arise from the three amino acid differences between the globular domains of H3.3 and H3.2 that mediate interaction with histone chaperones rather than from phosphorylation of S31. Although they develop normally, *H3.3^S31A^* adult males are infertile, consistent with a report of germ cell loss in *Drosophila* testes expressing H3.3S31A mutant histone (Chandrasekhara et al. 2023). Our observations differ from two previous studies; one that showed ectopic expression of H3.3AS31A rescues fertility in H3.3^Δ^ animals, and another that an *H3.3B^H3.2^* transgene rescues fertility of *H3.3^Δ^* animals (Hödl and Basler 2012; Sakai et al. 2009). Because transgenic *H3.3^S31A^*expression restores male fertility whereas expression from the endogenous genes does not, the functional defects of *H3.3^S31A^* might be compensated for by increased or modified gene expression. In females, H3.3—but not S31—is required for fertility, suggesting again that the AIG motif in the globular domain of H3.3 plays a critical role in H3.3 specific functions. Our studies do not rule out the possibility that H3.3S31 phosphorylation mediates certain stress responses, including signaling necessary for innate immunity.

### The genome is robust to changes in replication-independent H3 subtype identity

The behavioral defects of *H3.3^H3.2^* mutants prompted us to examine chromatin accessibility and gene expression profiles in the brain, which we found remained largely unaffected by loss of H3.3. This finding is consistent with previous studies showing that H3.3 depletion has only modest effects on gene expression and chromatin accessibility in mESCs (Banaszynski et al. 2013; Tafessu et al. 2023). Chromatin accessibility was similarly unchanged upon H3.3 deletion from mouse neural progenitor cells and in post-replicative neurons, although more extensive gene expression changes were reported in this study (Funk et al. 2022). The lack of chromatin accessibility defects in *H3.3* mutants is hard to reconcile with the requirement of H3 for nucleosome assembly and with the increase in H3.3 to near saturating levels during aging (Maze et al. 2015). However, our finding that replication-dependent H3.2 is normally expressed in adult brains offers a potential explanation, as it raises the possibility that H3.2 partially compensates for the absence of H3.3. Supporting this interpretation, we observed upregulation of replication-dependent H3.2 transcripts in *H3.3^Δ^* mutants, consistent with a prior observation (Sakai et al. 2009). These results suggest a homeostatic response that maintains functional levels of H3 in chromatin. We also observed increased H3.2 protein levels in *H3.3^Δ^* mutants; although this increase did not meet our strict statistical cutoffs, it is worth noting that small fold changes in histone abundance corresponds to hundreds of thousands of protein molecules due to the sheer abundance of histones in chromatin. To test this model of histone homeostasis, we depleted H3.2 via shRNA mediated knockdown in post-mitotic neurons and found that H3.2 is required in *H3.3^Δ^* mutants. Together, these findings support a model in which H3.2 expression contributes to a homeostatic mechanism that maintains an H3 amount sufficient to compensate for the absence of *H3.3*.

### Histone chaperones buffer against changes in histone supply

The observation that chromatin accessibility, gene expression, and developmental processes remain largely unaffected in *H3.2^H3.2^* mutants indicates that the histone supply-chain network can accommodate alterations in H3 subtype identity or expression. Indeed, most of the phenotypes we observed were linked to the H3 globular domain, suggesting that histone chaperones play a key role in the compensatory response. Structural and functional studies have identified the mechanisms by which chaperone complexes recognize and engage H3-H4, leading to the current model of “H3.2-specific” and “H3.3-specific” chaperones (Bhatt et al. 2025; Sitbon et al. 2020). For example, DAXX preferentially binds H3.3 via glycine 90 (H3.3G90), and when H3.3G90 is mutated to the H3.2-specific methionine (H3.3G90M), interaction with DAXX decreases and interaction with CAF-1 increases (Elsässer et al. 2012). Similarly, the UBN1/UBN2 subunits of HIRA interact via the A89 and G90 residues of H3.3, and these residues are necessary and sufficient for H3.3-specific recognition and binding (Xiong et al. 2018). Several studies suggest that these binding specificities underlie functional differences between replication-dependent H3 and replication-independent H3.3 *in vivo*. For example, in *Drosophila* Kc cells, ectopically expressed H3.2-GFP is not incorporated into chromatin in a replication-independent manner unless the globular domain residues are mutated to their H3.3 identities (Ahmad and Henikoff 2002b). Over-expression of replication-dependent H3.1 in neural stem cells lacking H3.3 does not rescue neural progenitor cell proliferation defects and over-expression of H3.2 in H3.3-depleted mESCs does not rescue defects in transcription of lineage-specific transcription factors (Xia and Jiao 2017; Banaszynski et al. 2013). How can these results be reconciled with our data indicating that H3.2 can functionally substitute for H3.3?

Intriguingly, several studies have reported that histone-chaperone interactions are more flexible *in vivo* than previously thought. For instance, CAF-1 associates with both H3.2 and H3.3 in HEK293T cells (Siddaway et al. 2022), and loss of DAXX in mouse fibroblasts leads to H3.3 association with CAF-1 (Drané et al. 2010). Our findings build on the emerging view of flexibility within the histone-chaperone network by suggesting that in the absence of H3.3 the *Drosophila* HIRA chaperone complex utilizes H3.2 to maintain genome and organism function. This model would explain why *Hira* becomes essential when replication-independent H3.3 is absent. We propose that histone chaperones can respond to changes in histone availability and engage alternative histone subtypes when the network is perturbed. However, histone chaperones likely interact less efficiently with non-cognate histones, causing flexibility of the histone chaperone network to have limits. Supporting this interpretation, we find that *H3.3^H3.2^* only partially compensates for the loss of *H3.3* in adult locomotor behavior and in longevity. We also find that the histone chaperone network is highly sensitive to gene dosage. Reducing *yem* (*UBN1*) dosage in animals with decreased replication-dependent *H3.2* and replication-independent *H3.3* gene copy number causes reduced viability (McPherson et al. 2023). Likewise, we find here that reducing replication-dependent histone gene dosage in *H3.3^H3.2^* animals causes complete lethality. These findings suggest that the compensatory capability within the histone-chaperone network is limited, and exceeding a minimum threshold of histone or chaperone abundance results in loss of chromatin homeostasis and developmental failure. For similar reasons, exceeding a maximum threshold may also result in loss of homeostasis, potentially explaining why some studies found that over-expression of replication-dependent histone H3 cannot rescue H3.3 deletion.

Altogether, by leveraging precise genome engineering, we demonstrate that the genome and development is robust to changes in the composition of the pool of replication-independent H3. We uncover a role for the HIRA histone chaperone complex in buffering against loss of replication-independent H3.3 to preserve genome function. These findings illuminate the dynamic interplay between histones and their chaperones in safeguarding chromatin integrity and organismal fitness. As histone dysregulation is increasingly implicated in aging and disease, our work provides a foundational framework for understanding how cells adapt to disruptions in histone supply.

## MATERIALS AND METHODS

### Drosophila strains and husbandry

Detailed fly strains and origins are described in **Supplemental Methods.** Fly stocks were maintained at 25°C on standard corn medium provided by Archon Scientific (Durham, NC) and LabExpress (Ann Arbor, MI).

### Generation of H3.3^Δ^, H3.3^H3.2^, and H3.3^S31A^ mutant lines

gRNA sequences were selected using FlyCRISPR Target Finder and CRISPOR (Concordet and Haeussler 2018; Gratz et al. 2015; Iseli et al. 2007), and cloned into pCFD4 (flycrisprdesign.org) along with protospacer sequences (Port et al. 2014). Embryo injections were performed by *GenetiVision* (Houston, TX). Details regarding CRISPR designs and diagnostic screening can be found in **Supplemental Methods**.

H3.3B^H3.2^ and H3.3B^S31A^: Two gRNA sequences targeting *H3.3B* were inserted into pCFD4 and co-injected into *yw; nos-Cas9(y+)/CyO; +/+* (Iso 3-19) with a 472bp single-stranded DNA repair template encoding the four H3.3B^H3.2^ substitutions (S31A, A87S, I89G, G90M) plus silent point mutations in the gRNA PAM sequences. *H3.3B^H3.2^*and *H3.3B^S31A^* alleles were identified using PCR of genomic DNA followed by BccI digestion. Three independent *H3.3B^H3.2^*and one *H3.3B^S31A^* mutant lines were isolated. His3.3B reference sequence: NCBI Gene ID 31848.

H3.3A^S31A^ and *H3.3A^Δ^*: Two gRNAs targeting *H3.3A* were inserted into pCFD4 and co-injected with a 461bp single-stranded DNA repair template encoding the four H3.3A^H3.2^ substitutions (S31A, A87S, I89V, G90M) plus silent point mutations in the PAM sites (*y1 M{nos-Cas9.P}ZH-2A w\**;; RRID:BDSC_54590). *H3.3A^S31A^* alleles were identified using PCR of genomic DNA followed by BccI and EagI digestion. One *H3.3A^S31A^* mutant line was isolated. Multiple *H3.3A^Δ^*alleles were identified using PCR of genomic DNA. H3.3A^H3.2^: Two gRNAs targeting *H3.3A* were inserted into pCFD4 and co-injected with a circular 2.25-kb homologous repair template encoding the four H3.3A^H3.2^ substitutions (S31A, A87S, I89G, G90M) plus silent point mutations in the gRNA PAM and seed sequences (*y1 M{nos-Cas9.P}ZH-2A w\**; RRID: BDSC_54590). *H3.3A^H3.2^* alleles were identified using sanger sequencing of genomic DNA. PCR primers targeting the pUC19 repair template backbone were used to confirm the absence of repair template backbone integration. One *H3.3A^H3.2^* mutant line was isolated. His3.3A reference sequence: NCBI Gene ID 33736. For each CRISPR-edited strain used, the *H3.3* gene was sequenced to verify the absence of additional mutations.

### Generation of H3.3B^WT^ transgene

*H3.3B* coding and regulatory regions (2363bp upstream of start codon to 2305bp downstream of stop codon were inserted into pJFRC177-10XUAS-FRT>-dSTOP-FRT>-myr::GFP (Gerald Rubin) using the NEB HiFi kit (New England Bio Labs, E5520S). The construct was inserted into ZH-86Fb (chr3R 86F8). See **Supplemental Methods** for primer sequences.

### Reverse transcription PCR

RNA was isolated from 3-5 adult males using the Zymo Direct-zol RNA Miniprep Kit (Zymo Research). cDNA was synthesized using the SuperScript III First-Strand Synthesis System (Invitrogen, random hexamers). *H3.3B^0^*, *H3.3B^H3.2^* and *H3.3B^S31A^*PCR products were digested with BccI. *H3.3A^Δ^*, *H3.3A^H3.2^* and *H3.3A^S31A^* PCR products were digested with BccI and EagI. Uncut and cut samples were run on polyacrylamide gels (8% in 1xTBE) or agarose gels (1%). Primers and screening parameters can be found in **Supplemental Methods.**

### Viability

Vials were maintained at 25°C and flipped every other day. Data for three biological replicates were plotted as the percentage of expected that were observed based on predicted Mendelian ratios for each cross. Chi square analysis was performed to determine statistical significance (p<0.05). Raw data can be found in: **Supplemental Table S1** (Fig. 1C), **Supplemental Table S12** (Fig. 6A), **Supplemental Table S13** (Fig. 6B), and **Supplemental Table S14** (Fig. 6C). See **Supplemental Methods** for respective crossing schemes. All genotypes were confirmed by PCR.

### Fertility

Single male and virgin female flies were transferred to vials with 3 wild-type adults of the opposite sex, and after 10 days the vials were examined for eggs that hatched into larvae. 10 individual animals were used in each biological replicate. Data (**Supplemental Table S2)** were plotted as mean ±SEM. Significance was determined by one-way ANOVA with post-hoc Tukey HSD.

### Negative geotaxis assay

Eight to twelve one– or 10-day old flies were separated by sex and placed in two empty vials connected face to face with white tape. After 10 minutes of acclimation, testing was performed by tapping down the flies three times in quick succession and determining the percentage of flies that climbed past a mark 8 cm from the bottom of the chamber within 30 seconds. Three biological replicates (10 trials per replicate) were performed for each genotype, and results were plotted as mean ±SEM. Significance was determined by one-way ANOVA with post-hoc Tukey HSD. See **Supplemental Tables S3** and **Supplemental Table S15** for more details.

### Longevity assay

Batches of 10 virgin females were transferred to fresh vials every Monday, Wednesday, and Friday, and the number of live flies was scored until all animals died. Dead flies were removed from vials. Ten batches were assayed per genotype. Survivorship curve was calculated using a Kaplan-Meier approach. Pairwise survival comparisons were performed using the log-rank test and resulting p values were adjusted for multiple testing using the Benjamini-Hochberg procedure (**Supplemental Table S4)**.

### RNA-sequencing library preparation

8 brains were dissected per replicate from 1d and 10d old flies in ice cold 1X PBS and solubilized in ice cold Trizol Reagent (Invitrogen #15596026). RNA was isolated using the Zymo Direct-zol RNA Miniprep kit (Zymo Research #ZR2070) including treatment with DNAse I. Libraries were prepared from total RNA using the Tecan Universal Plus Total RNA-seq with NuQuant kit, with custom *Drosophila melanogaster* rRNA depletion (Tecan #30185152) and sequenced at the UNC High Throughout Sequencing Facility using the NextSeq 2000 paired-end 100 platform.

### RNA-sequencing analysis

Sequence reads were trimmed for adaptor sequence/low-quality sequence using BBDuk (bbmap v38.67) with parameters: ktrim=r, k=23, rcomp=t, tbo=t, tpe=t, hdist=1, mink=11. Reads were mapped to a custom dm6 genome where the *HisC* region (chr2L:21,400,839-21,573,418) was deleted and a 1x Histone array (5kb repeat) was included as a new chromosome (chrHis). The modified genome file can be found at GEO:GSE317848. Genome files for use with the STAR aligner were generated using parameter: sjdbOverhang 99. Paired-end sequencing reads were aligned using STAR v2.7.7a with default parameters (Dobin et al. 2013). featureCounts (subread v2.0.6) was used to count reads mapping to features with parameters: –t exon, –s 1, ––countReadPairs (Liao et al. 2014). DESeq2 (v1.34.0) was used to identify differentially expressed genes (Love et al. 2014). All samples and replicates were included in the DESeq2 counts matrix. Differentially expressed genes were defined as genes with an adjusted p value less than 0.05 and an absolute log2 fold change greater than 1 after using ‘ashr’ for logFoldChange shrinkage (Stephens 2017). Principle component analysis and Pearson correlation were used to assess biological replicate similarity (**Supplemental Fig. S4).** Gene set enrichment analysis was completed using gseGO in clusterProfiler with parameters: ont = BP, minGSSize = 25, maxGSSize = 400, pvalueCutoff = 0.05, pAdjustMethod = BH (Wu et al. 2021; Yu et al. 2012). RPGC normalized bigwig files were used for data visualization in IGV (Robinson et al., 2011). Volcano plots, GSEA plots, and heatmaps were generating using ggplot2 (Wickham 2016). See **Supplemental Tables S8,S9** for DESeq2 results and GEO:GSE317848 for featureCounts output and pooled bigwig files. For analysis of replication-dependent histone transcript levels the gtf file used for featureCounts was modified to include an extended 3’ UTR for each replication-dependent histone gene to count reads mapping downstream of the normal replication-dependent histone mRNA 3’ ends. DESeq2 was run as described above. DESeq2 results can be found in **Supplemental Table S10**.

### ATAC-sequencing library preparation

3 brains were dissected per replicate from 1d and 10d old flies and cell dissociation, nuclear isolation, transposition, and DNA purification were performed as described in Buchert et al. 2023. Transposed DNA was amplified using the NEBNext High-Fidelity 2x PCR Master Mix (NEB, M0541S) and Nextera barcoded PCR primers. Library PCR amplification conditions can be found in Supplemental Methods.

### ATAC-sequencing analysis

Raw fastq files were processed using the nf-core atacseq v2.1.2 pipeline using Nextflow v24.04.2 (Di Tommaso et al. 2017). Reads were aligned to the dm6 genome using bowtie2. Reads mapping to chrM and blacklist regions were removed. Peaks were called using MACS2 with a p value cutoff of 0.05 (Zhang et al. 2008). High confidence peaks were then identified using the Irreproducible Discovery Rate (IDR) framework with a p value cutoff of 0.05 (Li et al. 2011). Change in accessibility across high-confidence peaks was assessed using DESeq2 (Love et al. 2014). Differentially accessible peaks comparing 1d vs 10d were defined as peaks with an adjusted p value less than 0.05 and an absolute log2 fold change greater than 0.585 after using ‘ashr’ for logFoldChange shrinkage. Differentially accessible peaks comparing different genotypes at a single time (1d or 10d) were defined as peaks with an adjusted p value less than 0.05 and an absolute log2 fold change greater than 1 after using ‘ashr’ for logFoldChange shrinkage. Principle component analysis was used to assess biological replicate similarity (**Supplemental Fig. S4).** RPGC normalized bigwig files were used for data visualization in IGV and for average signal plots generation using deepTools2 (Ramírez et al. 2016; Robinson et al. 2011). Volcano plots, GSEA plots, and heatmaps were generating using ggplot2 (Wickham 2016). See GSE317810 for peak calls and pooled bigwig files.

### Western blotting

Protein extracted from *H3.3^WT^, H3.3^Δ^, H3.3^H3.2^, and H3.3^S31A^* 1d heads was subjected to western blotting as described in McPherson et al. 2023 using the following antibodies: rabbit anti-H3 (1:10,000; Abcam Cat# ab1791, RRID:AB_302613), rabbit anti-H3.3 (1:1,000, Abcam Cat# ab176840, RRID:AB_2715502), rabbit anti-H3.1/H3.2 (1:1,000, Millipore Cat# ABE154, RRID:AB_2811170), and donkey anti-Rabbit-IgG-HRP (1:10,000, Cytiva Cat# NA934, RRID:AB_772206) secondary antibody. Blots were detected using Amersham ECL Prime Western blotting Detection Reagent (Cytvia, RPN2232). ImageLab densitometry analysis was used to determine H3.2, H3.3, and H3 band intensity. Histone signal was normalized to corresponding total protein signal. Normalized signals from different titrations of the same genotype were averaged and resulting values were set relative to the wild-type value. This process was completed for three biological replicates. See **Supplemental Table S11** for statistical analysis, replicate signal values, and replicate images.

### Use of generative AI

During the preparation of this work, the author(s) used ChatGPT and GitHub Copilot to edit and generate code for data visualization. After using these tools, the authors reviewed and edited the content as needed and take full responsibility for the content of the publication.

### Data availability

High throughput sequencing data sets have been deposited at Gene Expression Omnibus (GEO) as GSE317848 and GSE317810 and are publicly available as of the date of publication.

## COMPETING INTEREST STATEMENT

The authors declare no competing interests.

## ACKNOWLEDGMENTS

We gratefully acknowledge the technical support from the UNC High Throughput Sequencing Facility. This work was supported by NIGMS grants R35GM128851 to D.J.M., R35GM145258 to R.J.D. J.E.M. was supported by NIGMS grant T32GM135128 and NIA grant F31AG079632.

## AUTHOR CONTRIBUTIONS

Conceptualization, J.E.M, D.J.M, and R.J.D; methodology, J.E.M, D.J.M, and R.J.D; Investigation, J.E.M, C.A.H, C.S, M.P.L.J, and L.C.G; writing—original draft, J.E.M; writing—review & editing, J.E.M, R.J.D and D.J.M; funding acquisition, J.E.M, R.J.D, and D.J.M; supervision, D.J.M and R.J.D.

